# First Crystal Structure of a Non-Structural Hepatitis E Viral Protein Identifies a Putative Novel Zinc-Binding Protein

**DOI:** 10.1101/538884

**Authors:** Andrew Proudfoot, Anastasia Hyrina, Meghan Holdorf, Andreas O. Frank, Dirksen Bussiere

## Abstract

Hepatitis E virus (HEV) is a 7.2 kb positive-sense, single-stranded RNA virus containing three partially overlapping reading frames, ORF 1-3. All non-structural proteins required for viral replication are encoded by ORF1 and are transcribed as a single transcript. Computational analysis of the complete ORF1 polyprotein identified a previously uncharacterized region of predicted secondary structure bordered by two disordered regions coinciding partially with a region predicted as a putative cysteine protease. Following successful cloning, expression and purification of this region, the crystal structure of the identified protein was determined and identified to have considerable structural homology to a fatty acid binding domain. Further analysis of the structure revealed a metal binding site, shown unambiguously to specifically bind zinc via a non-classical, potentially catalytic zinc-binding motif. We present analysis for the first time of this identified non-structural protein, expanding the knowledge and understanding of the complex mechanisms of HEV biology.

**Importance:** Hepatitis E virus (HEV) is an emerging virus found predominately in developing countries causing an estimated 20 million infections, which result in approximately 57,000 deaths a year. Although it is known that the non-structural proteins of the HEV ORF1 are expressed as a single transcript, there is debate as to whether ORF1 functions as a single polyprotein or if it is processed into separate domains via a viral or endogenous cellular protease. In the following paper, we present the first structural and biophysical characterization of a HEV non-structural protein using a construct that has partially overlapping boundaries with the predicted putative cysteine protease. Based on the structural homology of the HEV protein with known structures, along with the presence of a catalytic zinc-binding motif, it is possible that the identified protein corresponds to the HEV protease, which could require activation or repression through the binding of a fatty acid. This represents a significant step forward in the characterization and the understanding of the molecular mechanisms of the HEV genome.

## Introduction

Hepatitis E virus (HEV) is a non-enveloped, positive single-stranded RNA virus that infects approximately 20 million people annually, with a global mortality rate of 2% (1, 2). This fatality rate dramatically increases up to 30 % for infected pregnant women in their third trimester for unknown reasons (3). The virus is transmitted via the fecal-oral route and is most common in semi-tropical and developing countries with poor sanitation or contaminated water supplies (4), however cases of HEV infection reported in more industrialized countries in Europe, as well as in the USA and in Japan are increasingly more common (5–7). Although currently there is no accepted treatment for HEV infection, the off-label treatments of both interferon and/or ribavirin have been used successfully to treat chronic HEV infection (8, 9). As HEV is a growing worldwide threat, better understandings of the viral molecular mechanisms are necessary to help enable the development of targeted therapeutics.

HEV is a member of the *Hepeviridae* family comprised of four main mammalian genotypes; genotypes 1 and 2 are infection limited to only the human host, while genotypes 3 and 4 are zoonotic and have been identified in humans along with other animal species including swine (10). The HEV genome is approximately 7.2 kb in size and is comprised of three partially overlapping open reading frames (ORF) flanked by 5’ and 3’ untranslated / non-coding regions (11–13). ORF1 encodes all the non-structural proteins required for viral replication. ORF2 is the only ORF that has been structurally characterized in the HEV genome, and encodes the viral capsid protein that is involved with virion assembly (13). ORF3, the smallest of the three ORFs, encodes a 113 - 115 amino acid protein implicated to either encode an ion channel or aid in HEV virion release (14). In addition, the HEV genome contains a 7-methylguanosine cap at the 5’ end, and two *cis*-reactive elements that are dispersed through the viral genome, all of which are essential for viral replication (15, 16).

The largest of the three ORFs is ORF1, which is transcribed as a single 1693 residue polyprotein (17). In the absence of structural characterization, ORF1 has been compared to homologous viruses and computationally analyzed to identify eight putative domains (18). These include a methyltransferase domain (Met), Y domain (Y), papain-like cysteine protease (PCP), a proline-rich region that contains a hypervariable region (H), X-domain (X), helicase (Hel) and an RNA dependent RNA polymerase (RdRP) (Figure 1). With the exception of the Y-domain and papain-like cysteine protease, all identified domains have now been characterized to some extent, and experiments have been performed to confirm the predicted protein functions (19–21). The predicted protease is more elusive as a distinct protease is not identifiable in the HEV polypeptide sequence, however homology modelling of other alphaviruses homologous to HEV which contain a cysteine protease, predict a putative cysteine protease between residues 433 and 592 in ORF1 (18).

**Figure 1:**
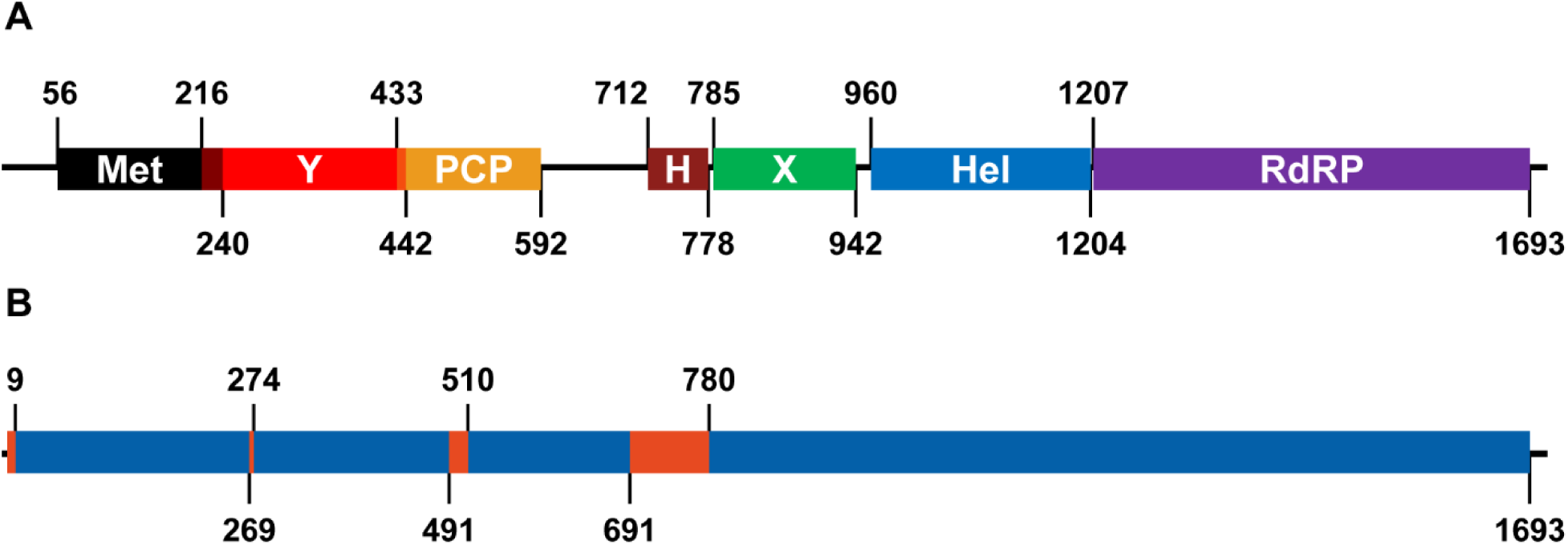
Schematic representations of the HEV ORF1 depicting the boundaries of the A) methyltransferase (Met), Y-domain (Y), putative cysteine protease (PCP), disordered hinge region (H), X-domain (X), helicase (Hel) and RNA dependent RNA polymerase (RdRP) as identified by Koonin *et al.* (18) and B) the predicted disordered (Orange) and secondary structure containing (Blue) regions.

Previous research has been unable to confirm the processing of ORF1 or the presence of a protease. Some reports show that ORF1 is expressed as a single 186 kDa polyprotein in *E. coli*, insect and mammalian cells, and in a cell free system (22). Although other reports support polyprotein processing of ORF1, when expressed in mammalian cells two peptides of approximately 107 and 78 kDa in size are observed (23). Likewise, expression of ORF1 in insect cells results in eight small peptides which can be inhibited in the presence of a cell-permeable cysteine protease inhibitor (24). Current research on the purified putative protease domain *in vitro* has been limited. Protein purified under denaturing conditions has been shown upon refolding to have proteolytic activity against ORF1 and ORF2 products, and mutation analysis of highly conserved cysteine and histidine residues in the predicted protease inhibit proteolytic activity and are essential for viral replication (25). A three-dimensional model of the predicted protease region has been produced using consensus-prediction results of the closely related p150 rubella protease (26), however characterization of the predicted protease domain is required in order to understand better the viral biology. In this study, we report a comprehensive analysis of the biochemical, biophysical and structural parameters of the predicted protease domain that may help elucidate the function of the protein in the HEV lifecycle.

## Results

All previous work performed on the HEV non-structural proteins of ORF1 has been done based on the boundaries originally proposed by Koonin *et al* (18) (Figure 1A), with the resulting protein constructs typically produced with low yields or requiring refolding from the insoluble fraction following purification (11, 20, 25). To evaluate if these difficulties were due to distinct differences between the actual and proposed domain boundaries, ORF1 was analyzed using secondary structure prediction to identify regions of order and disorder within the open reading frame (Figure 1B). The HEV X-domain, helicase and RNA-dependent RNA polymerase show high levels of homology to other homologous proteins, so it was not surprising that secondary structure is predicted between residues 780 – 1693 in ORF1. Our analysis also showed that secondary structure elements at the N-terminus of ORF1 are sandwiched between four regions of disorder within ORF1, residues 1 – 8, 270 – 273, 492 – 509 and 692 – 779. The region of predicted disorder between residues 692 – 779 aligns with the region previously identified to be the disordered hinge region (residues 712 – 778), with the three remaining regions of disorder bordering three regions of predicted secondary structure. The two regions of order identified between residues 9 – 269 and 274 – 491 coincide with the previously identified methyltransferase and Y-domain respectively (18), but surprisingly our analysis revealed a previously unidentified predicted domain between residues 510 – 691. Prior analysis of this region had shown residues 433 – 592 contained the putative cysteine protease, however the prediction used to locate the putative protease was not conclusive and was based on two assumptions: HEV contains a cysteine protease and the proposed catalytic residues align with the identified distant homologous proteases (18). In addition, subsequent mutagenesis studies and homology modeling of this region have been performed to support the presence of the protease, but a protein structure and elucidation of the exact catalytic mechanism is required to fully understand any potential HEV protease (25–27). To confirm the presence of a domain between residues 510 and 691, a construct containing this region was codon-optimized, cloned and expressed in *E. coli* as an MBP-fusion protein (HEV^510-691^). Following cleavage of the fusion protein and purification, a final yield of approximately 120 mg of soluble protein per liter of unlabeled medium was achieved, and analysis by 1D NMR confirmed that the protein was folded. This high-level of expression suggests that this protein is extremely stable.

The protein was concentrated to 10 mg/ml and 864 crystallization conditions were screened at two temperatures, 4 °C and 20 °C. Two very similar conditions, which differed only by the type of nitrate present, provided initial hits (Figure 2A). Optimized crystals grew at 4 °C in precipitant consisting of 18 % w/v PEG 3350, 180 mM LiNO_3_ and 10 mM NiCl_2_ (Figure 2B) and diffracted to 1.8 Å resolution on an in-house X-ray source. Analysis of the RCSB showed that HEV^510-691^ did not have sequence homology to any deposited structure, so molecular replacement was not possible. Given this, the structure was solved using a mercury heavy-atom derivative and single-isomorphous replacement with anomalous scattering (SIRAS) (Table 1). The structure consists of ten β-strands and four α-helices, with the β-strands 1 – 4 and 5 – 10 arranged in two antiparallel sheets arranged in a sandwich-like fold. α-helices 1 & 2 are located between β-strands 1 & 2 and α-helices 3 & 4 are positioned at the C-terminus of the protein, with α-helix 4 sitting between the two antiparallel β-sheets (Figure 2C).

**Figure 2:**
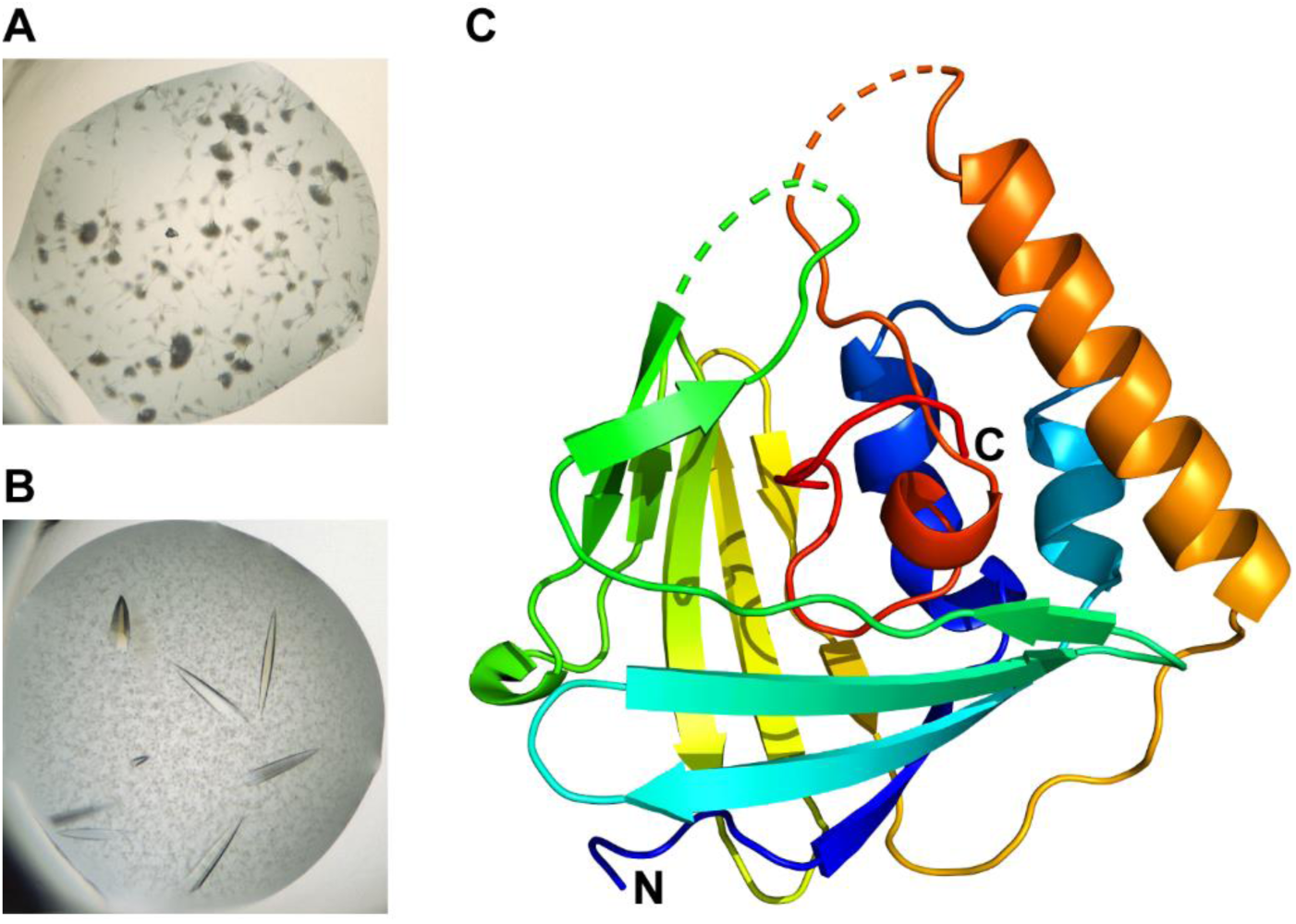
Crystallization and structure solution of HEV^510-691^. A) Initial crystal hits obtained in 18 % w/v PEG 3350, 180 mM LiNO_3_ as precipitant and grown at 4°C. B) Crystals obtained following optimization of the initial screening hit with 10 mM NiCl_2_. C) Structure of HEV^510-691^ shown in rainbow color representation with the N-terminus shown in blue and the C-terminus shown in red (PDB 6NU9).

**Table 1:** Data collection and refinement statistics of the HEV^510-691^ X-ray structure.

| Coordinate Set | HEV<sup>510-691</sup> Native | HEV<sup>510-691</sup> Thiomersal<br>Derivative | HEV<sup>510-691</sup> Zinc<br>Co-Structure |
| --- | --- | --- | --- |
| **PDB ID** |  |  | 6NU9 |
| **Data Collection Statistics** |  |  |  |
| Resolution (Å) | 45.56–1.95 (2.02-1.95) | 45.54-2.10 (2.18-2.10) | 52.62–1.76 (1.82–1.76) |
| Space group | P6<sub>5</sub>22 | P6<sub>5</sub>22 | P6<sub>5</sub>22 |
| Cell dimensions<br>a, b, c (Å) | 64.49, 64.49, 157.41 | 64.20, 64.20, 158.70 | 64.40, 64.40, 158.90 |
| $\alpha$, $\beta$, $\gamma$ (°) | 90.00, 90.00, 120.00 | 90.00, 90.00, 120.00 | 90.00, 90.00, 120.00 |
| Total reflections | 891654 (69081) | 378180 (29740) | 680486 (19154) |
| Unique reflections | 14844 (1430) | 21378 (2114) | 20137 (1960) |
| Multiplicity | 60.1 (48.3) | 17.7 (14.1) | 33.8 (9.8) |
| Completeness (%) | 99.4 (99.2) | 99.9 (98.9) | 99.9 (99.3) |
| I/$\sigma$(I) | 56.6 (22.5) | 20.5 (8.1) | 13.4 (1.2) |
| R<sub>merge</sub> | 0.068 (0.218) | 0.101 (0.333) | 0.228 (1.028) |
| R<sub>pim</sub> | 0.009 (0.032) | 0.023 (0.090) | 0.038 (0.334) |
| CC1/2 | 0.999 (0.998) | 0.999 (0.975) | 0.997 (0.339) |
| **Phasing Statistics** |  |  |  |
| Resolution (Å) |  | 38.24-2.50 |  |
| Heavy atom sites |  | 3 Hg, 2 Ni |  |
| FOM |  | 0.400 |  |
| **Refinement Statistics** |  |  |  |
| R<sub>work</sub> |  |  | 0.236 |
| R<sub>free</sub> |  |  | 0.268 |
| Number of non-H atoms |  |  |  |
| protein |  |  | 1306 |
| ligands |  |  | 10 |
| solvent |  |  | 339 |
| Protein residues |  |  |  |
| RMS bonds(Å) |  |  | 0.004 |
| RMS angles (°) |  |  | 0.66 |
| Ramachandran favored (%) |  |  | 96.30 |
| Ramachandran allowed (%) |  |  | 3.70 |
| Ramachandran outliers (%) |  |  | 0 |
| Clashscore |  |  | 3.81 |
| Average B-factor ($\text{\AA}^2$) | | | 17.51 |

The crystal structure was analyzed using DALI to identify if there were any structurally homologous proteins in the PDB (28). The structural alignment showed that although there was only 13 – 15% amino acid conservation with the identified structural homologs, residues 514 – 635 of HEV^510-691^ has significant structural homology to multiple fatty acid binding domains, with the homologous regions aligning with a backbone RMSD of ∼ 2.5 Å (Figure 3A & B). This homologous region of the HEV protein corresponded to all β-strands and the first two α-helices, while helices 3 and 4 were not present in any of the structural homologs identified. In our structure, α-helix 4 is located between the anti-parallel β-sheets, and analysis of the identified homologous structures showed fatty acids bound in the same location (Figure 3C). An additional HEV construct from residues 510 – 635 (HEV^510-635^), which has boundaries homologous to the identified fatty acid domains was produced. However, in this case the removal of the C-terminal residues abolished all protein expression, indicating that the additional residues and resulting secondary structure were likely part of the identified domain and may play an important role in protein folding and/or function.

**Figure 3:**
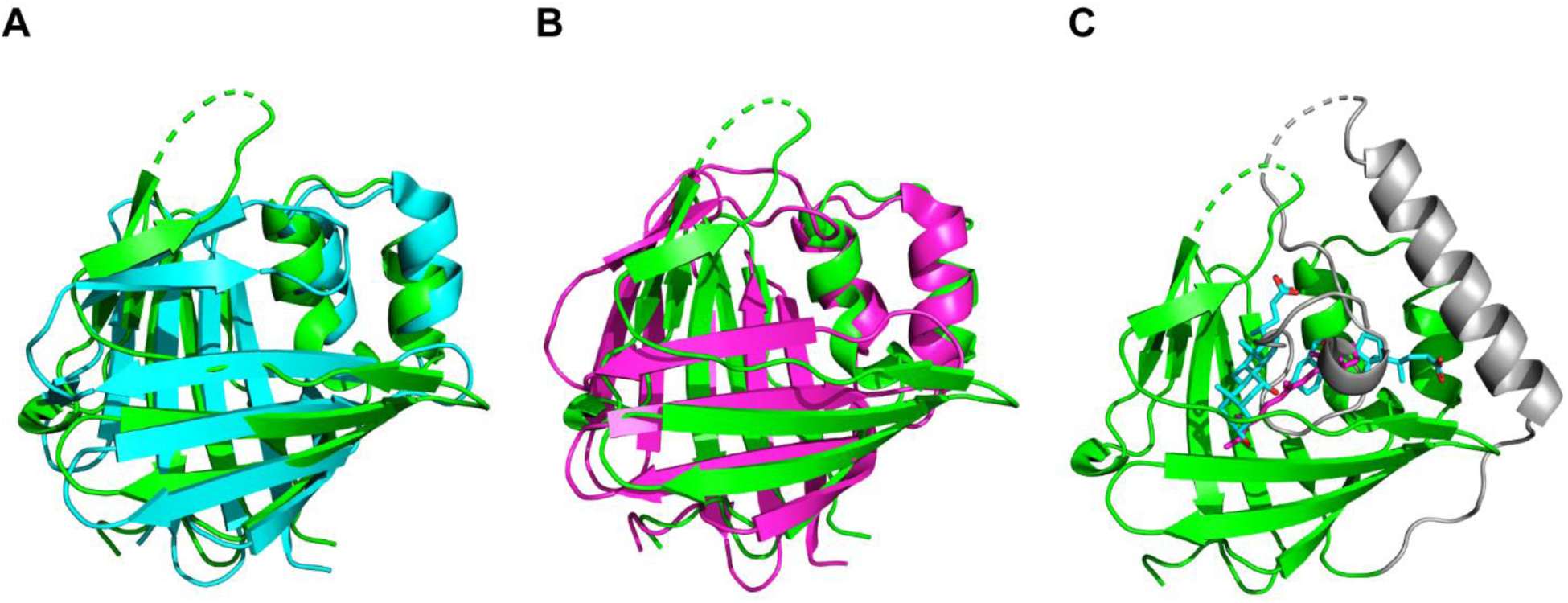
Superposition of HEV^510-691^ with the most significant structural homologs identified in DALI. Residues 514 – 635 of HEV^510-691^ with A) chicken liver fatty acid binding protein (49) (PDB 1TW4) Z-score 10.6 and B) retinol binding protein (PDB 5FEN) Z-score 11.2. C) Superposition of the bound cholic acid (cyan) present in 1TW4 and retinol (magenta) present in 5FEN with HEV^510-691^.

Analysis of the region around α-helices 3 and 4 identified that there was density corresponding to a bound metal ion, coordinated by two residues, His 671 and Glu 673. In addition, another residue, His 686, was positioned in close proximity to act potentially as the third co-ordination site. However, in our structure, His 686 was in the opposite conformation and the side chain was not oriented to coordinate the bound metal (Figure 4A). Native HEV^510-691^ crystals soaked in a 1 mM solution of EDTA showed signs of cracking following 1 hour of soaking and the resulting datasets showed no electron density in the region where we had previously identified the bound metal. This confirmed that the identified density corresponded to a metal or some other entity that can also be coordinated by EDTA. In the HEV^510-691^ crystal structure, the density corresponding to the metal atom could not be fully satisfied by a single metal during refinement. This indicated that there were either different metal species present (i.e. the endogenous metal in addition to metal from either the purification or the crystallization conditions), or that the binding site was not fully saturated with the endogenous metal and thus was partially occupied. To identify which endogenous metal the protein had highest affinity for, DSF was used to screen a number of different possibilities (Table 2) to see which, if any, metal could induce an increase in thermal stability. Of the 17 metals tested, zinc was the only metal that stabilized the protein and induced an increase in thermal stability of 3.0 °C (Figure 4B). Datasets acquired with HEV^510-691^ crystals soaked in a 1 mM solution of zinc chloride showed zinc at this position. However, the His 686 sidechain was still not orientated properly to act as the final zinc co-ordination site.

**Figure 4:**
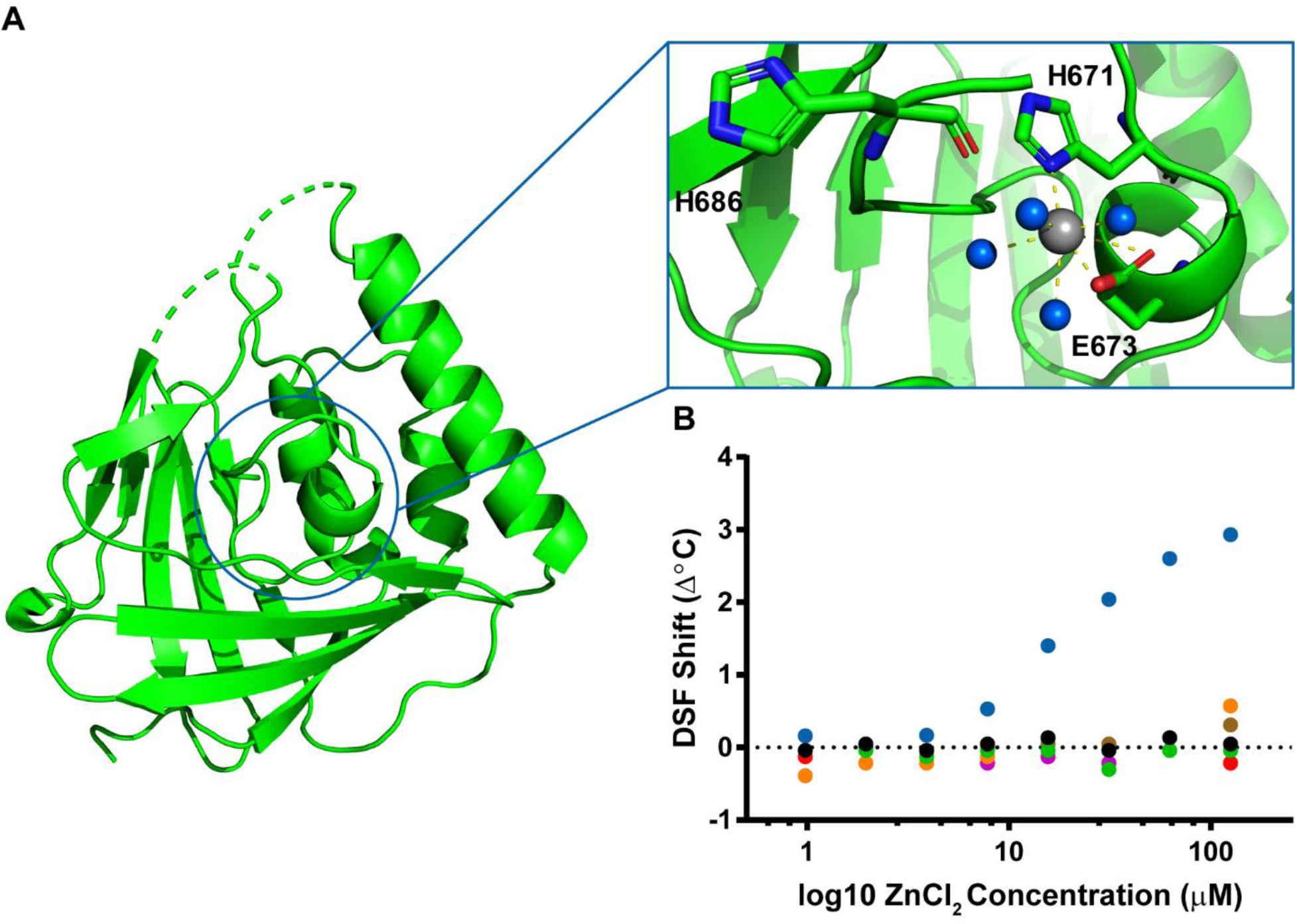
Analysis of metal interactions with HEV^510-691^. A) Magnification of region adjacent to the bound metal (grey) showing surrounding waters present (blue) and potential coordinating residues H671, E673 and H686. B) DSF analysis of wild type HEV^510-696^ in the presence of increasing concentrations of zinc (blue), calcium (green), cobalt (brown), iron (orange), manganese (red) and magnesium (magenta).

**Table 2:** Metals screened by DSF for binding to HEV^510-691^.

| Metals that have a positive impact on protein stabilization |  |  |  |
| --- | --- | --- | --- |
| Zinc |  |  |  |
| Metals that destabilize of do not impact protein stability |  |  |  |
| Barium | Cadmium | Cesium | Calcium |
| Chromium | Cobalt | Copper | Iron |
| Lithium | Magnesium | Manganese | Nickel |
| Potassium | Praseodymium | Strontium | Yttrium |

His 686 is located close to the end of the crystalized construct, so it is plausible that additional residues at the C-terminus of the construct could induce the formation of additional secondary structure and change the orientation of the His 686 side chain. To test this, additional constructs were designed selecting C-terminal boundaries based upon the presence of stretches of low complexity amino acid sequences (i.e. poly-glycine, -alanine, -serine or combinations thereof) with the additional consideration of not adding a significant number of disordered residues to the HEV^510-691^ construct. A construct from residues 510 – 720 (HEV^510-720^) was produced that was 29 residues longer than HEV^510-691^, and was analyzed by NMR. Comparison of HEV^510-720^ with the HEV^510-691^ showed that the additional residues had amide chemical shifts in the region of 7.5 – 8.5 ppm, which is indicative of disordered amino acids (Figure 5A), and coincided with the performed secondary structure prediction (Figure 1B). Analysis by DSF showed that in the presence of zinc, the HEV^510-720^ had a greater increase in thermal stability (5.5 °C) compared to the HEV^510-691^ (Figure 5C). To optimize the construct further, additional C-terminal truncations were made and biophysically characterized to identify a construct that both added the fewest disordered residues and induced an equivalent increase in protein thermal stability in the presence of zinc. A construct from residues 510 – 696 (HEV^510-696^) was identified to be the optimal construct and exhibited a 6 °Cincrease in thermal stability in the presence of zinc (Figure 5C), but did not contain an additional increase in disorder in the NMR spectrum (Figure 5B). HEV^510-696^ was used for crystal screening, but no crystals were observed with this construct in the presence or absence of zinc.

**Figure 5:**
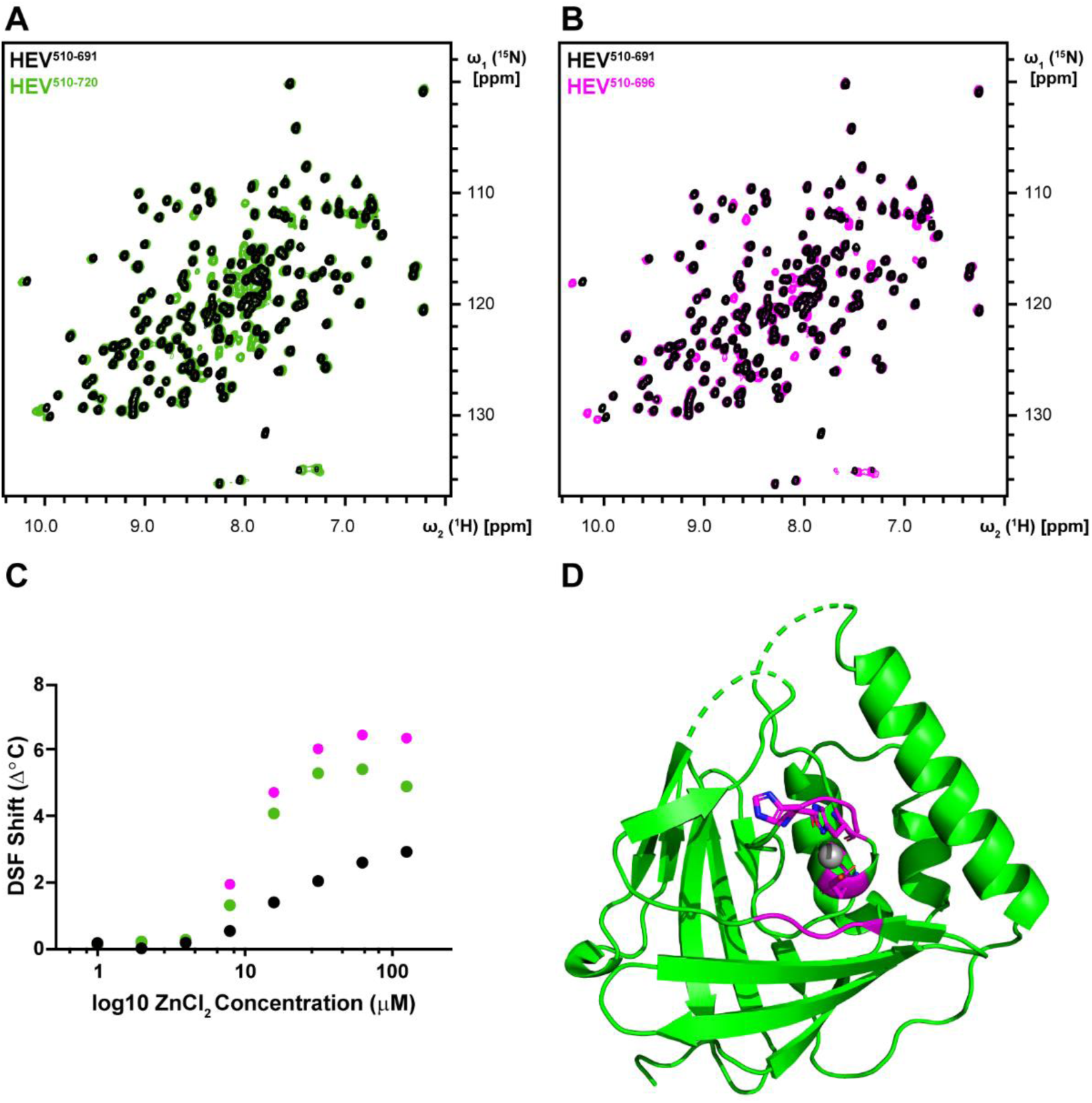
Optimization of the C-terminus of HEV^510-691^. Superposition of 2D [^15^N, ^1^H]-HSQC spectra acquired with HEV^510-691^ (black) and A) HEV^510-720^ (green) and B) HEV^510-696^ (magenta). C) DSF analysis of HEV^510-691^ (black), HEV^510-720^ (green) and HEV^510-696^ (magenta) in the presence of increasing concentrations of ZnCl_2_. Structure of HEV^510-691^ showing all residues (magenta) that experience a significant chemical shift perturbation when compared to HEV^510-696^.

When comparing the [^1^H – ^15^N] HSQC spectra of HEV^510-691^ and HEV^510-696^, although many of the signals in the two spectra overlapped, there were also a number of amide resonances that experienced a significant chemical shift perturbation. Assignments of all amide peaks were obtained, and the residues experiencing a significant chemical shift perturbation were mapped to the three potential zinc co-ordination residues, the residues at the C-terminus and a stretch of amino acids on β-strand 4, between residues 572 – 574 (Figure 5D). Although these constructs did not crystallize, it can be inferred from the NMR data that the additional C-terminal residues form a β-strand, running antiparallel to β-strand 4. This would also allow His 686 to be oriented in such a way as to act as the final zinc co-ordination site. To confirm if His 671, Glu 673 and His 686 were involved with zinc co-ordination, each of the three residues were mutated individually and together to an alanine residue. Mutation of each residue contributed approximately a 2 °C reduction in thermal stability when compared to HEV^510-696^(approximately 2 °C for each single mutation and 6 °Cfor the triple mutant). However in comparison to HEV^510-696^, all mutant constructs failed to show any increase in thermal stability in the presence of zinc (Figure 6), indicating that all three residues are involved with zinc co-ordination.

**Figure 6:**
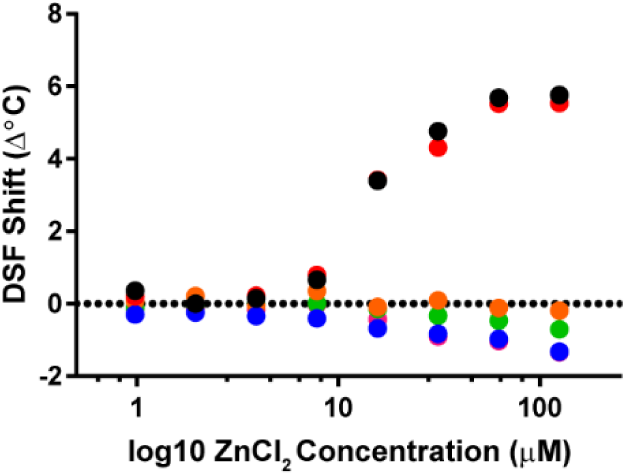
DSF analysis of HEV^510-696^ (black), ΔE583A (red), ΔH671A (blue), ΔE673A (orange), ΔH686A (green) and ΔH671A/ΔE673A/ΔH686A (magenta) HEV^510-696^ in the presence of increasing concentrations of ZnCl_2_.

Work characterizing the predicted HEV protease has been performed with constructs that closer matched the boundaries identified by Koonin *et al* (18, 25, 29) (Figure 1A). To investigate the effect of additional residues at the N-terminus of HEV^510-691^, additional constructs were produced that extended the N-terminal boundaries to residues 455 (HEV^455-691^), 440 (HEV^440-691^) and 403 (HEV^403-691^). These boundaries were selected as described above, in regions that had been predicted not to contain regular secondary structure. Previous work identified that *E. coli* BL21 cells expressing residues 440 – 610 experienced cell death upon induction at different temperatures and various concentrations of IPTG (25). Consistent with these observations, when the three N-terminally extended constructs were expressed in unlabeled medium in the absence of trace metals, cell growth was significantly inhibited following induction of the target proteins with IPTG. However, when *E. coli* were grown in unlabeled medium that had been supplemented with 1 x trace metals, normal cell growth and soluble protein expression was observed following induction with IPTG.

As the number of residues at the N-terminus increased, a corresponding reduction in protein expression was observed, with HEV^440-691^ identified to have comparable protein expression to the shorter constructs tested, which coincided with the previously identified putative protease domain (18). Following gel filtration of HEV^440-691^, it was apparent that the purified protein eluted from the gel filtration column at a much lower retention volume (Figure 7A) than expected (at a retention volume corresponding to a protein of Mw 270 kDa) and was brown in color. Mass spectrometry was performed with the protein species under denaturing conditions and a single peak at 27136.33 Da was observed (Figure 7B), which corresponded to the expected molecular weight of the monomeric protein and indicated that the protein eluting from the column was oligomerized. It was likely that the brown hue was coming from a bound metal and to identify if this was the case, the protein was thermally denatured, spun down and the supernatant was analyzed with 5F-BAPTA. 5F-BAPTA is divalent cation chelator with two fluorine atoms positioned at simultaneous positions, which yields a single peak in the ^19^F NMR spectrum. This signal can be monitored using ^19^F NMR as different bound metal ions induce different characteristic downfield chemical shift perturbations to the free 5F-BAPTA signal (30). Although a strong brown color was observed in the protein pellet following denaturation and centrifugation, the metal that remained in solution induced a 27.8 ppm chemical shift perturbation to the 5F-BAPTA signal (Figure 7C) which is consistent with the observed chemical shift upon 5F-BAPTA interacting with divalent iron (Fe^2+^) (28.1 ppm).

**Figure 7:**
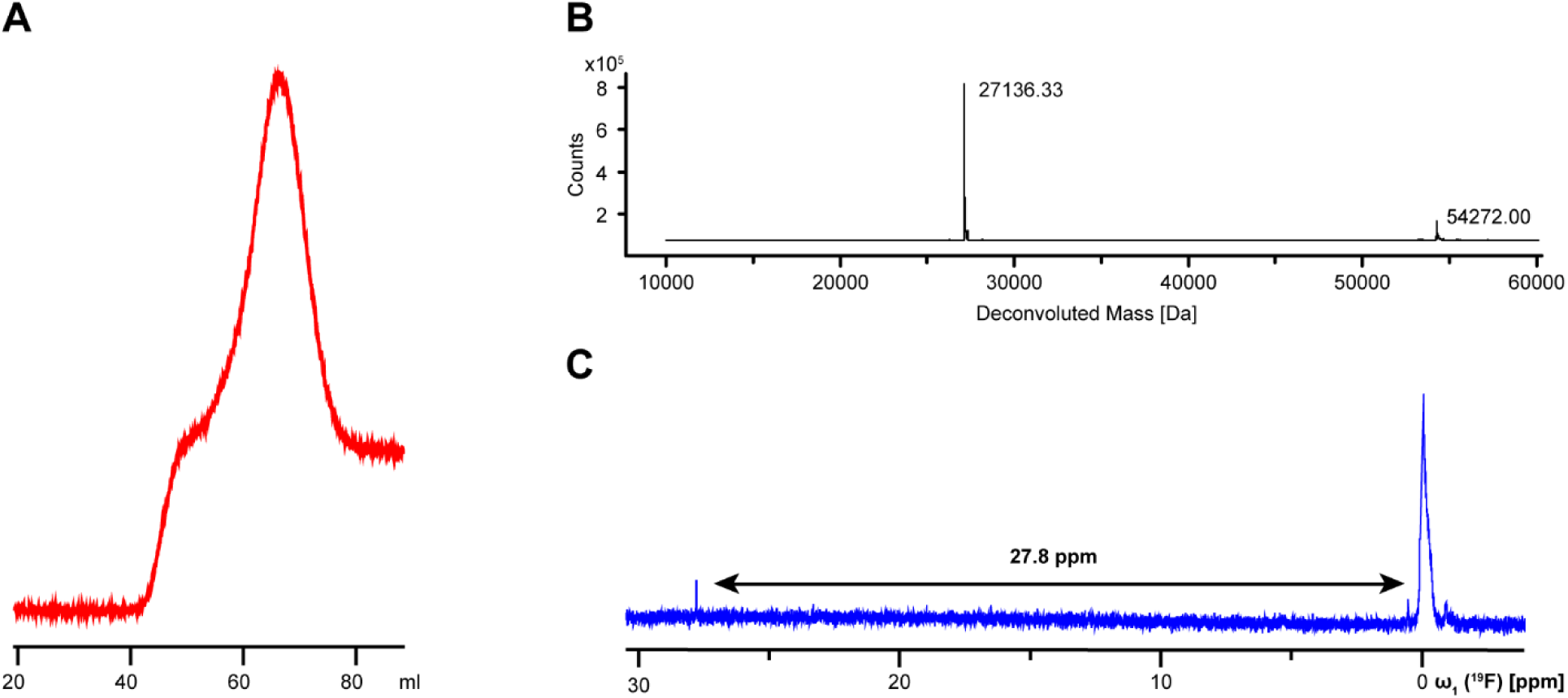
Purification and analysis of HEV^440-691^. A) Elution profile measured at 280 nm of HEV^440-691^ from a HiLoad 16/60 Superdex 75 gel filtration column. B) Mass spectrometry analysis of the purified HEV^440-691^. C) ^19^F 1D analysis of metals co-purified with HEV^440-691^, in complex with 5F-BAPTA.

## Discussion

Detailed characterization of the non-structural proteins in ORF1 has proven to be a significant impediment in the understanding of the HEV genome. Initial structural analysis found regions in the HEV ORF1 that were predicted to correspond to a methyltransferase, Y-domain, disordered hinge, X-domain, helicase and RNA dependent RNA polymerase. Based on other positive-sense RNA viruses, it was hypothesized that a protease should also be encoded within ORF1. However, structural analysis could not conclusively identify a corresponding region, and the proposed cysteine protease was assigned to a region between residues 433 and 592 (18).

Our analysis of the HEV ORF1 identified that there is a region of ordered secondary structure (resides 510 – 691) sandwiched between two regions of disorder (residues 492 – 509 and 692 – 779). Overexpression of this region resulted in a soluble, stable protein, suitable for crystallization. The resulting protein structure of the HEV^510-691^ displayed a high structural homology to fatty acid binding domains, that when compared to homologous fatty acid binding proteins, revealed two additional α-helices present at the C-terminus of the construct. One of the two additional α-helices is located between the two β-sheets, in the same position where homologous fatty acid binding proteins bind their respective fatty acids (Figure 3C). Removal of these two additional α-helices from the HEV construct resulted in no expression of the protein, suggesting that this region was important for structural stability.

Further analysis of the additional α-helices showed the presence of a bound zinc metal ion coordinated by residues His 671, Glu 673 and His 686, which was confirmed by DSF and site-directed mutagenesis studies (Table 2 and Figure 6). Each single mutant had a similar Tm in the absence of zinc (within 2°C) suggesting that each of these mutants abolished zinc coordination but did not adversely affect the protein fold. Although it is currently unknown if the coordinated zinc performs a catalytic or structural role, coordination of zinc for the purpose of structural integrity is predominantly achieved through a combination of cysteine and histidine residues, whereas acidic amino acids are included in the coordination site when the bound zinc is used for catalytic purposes (31). Analysis of all annotated HEV sequences present in Uniprot show complete conservation of residues Glu 673 and His 686, whereas residue 671 fluctuates between a histidine (78%) and a tyrosine (22%) in the available deposited sequences.

Taking into consideration the high conservation of the coordinating residues along with the observation that the protein has a potential catalytic zinc coordination site, it is possible that the solved structure corresponds to a zinc metalloprotease. All metalloproteases require an acidic residue in close proximity to the coordinated zinc to act as the catalytic residue during the proteolytic reaction. Analysis of the amino acid conservation of other residues in close proximity to the bound zinc identified the highly conserved Glu 583, which is positioned between β-strands 5 and 6, which could potentially serve as the catalytic residue. As expected, mutagenesis of Glu 583 to alanine did not have any significant effect on the induced increase in thermal stability upon addition of zinc as this residue is not directly involved with zinc coordination and thus does not directly stabilize the proteins tertiary structure. In the crystal structure of HEV^510-691^, the current geometries of the zinc coordinating residues are not optimal to catalyze a proteolytic reaction, however there remains the possibility that the binding of a fatty acid or some other endogenous ligand between the two β-sheets could displace α-helices 3 and 4, re-orientating the zinc coordinating amino acids into a catalytically active orientation. While we have not identified the endogenous ligand that interacts with the domain, it is well documented that the liver is rich in fatty acids so binding of a specific fatty acid to the protease could be used by the virus as a regulatory mechanism. In addition, it is known that the biochemical profile of the liver changes throughout pregnancy (32). Thus, if the protein has a specificity for a fatty acid that is at elevated levels in pregnant women, this could also explain why the virus has increased mortality and morbidity in pregnant women in their third trimester (3).

There are no reports in the literature either unambiguously confirming or refuting the existence of a protease in HEV, with assays performed *in vitro* and in different cell lines reporting several distinct cleavage products (22–25, 27, 33, 34). If the HEV protease requires binding of a particular fatty acid to correctly form the catalytic site for ORF1 processing, this could potentially explain these results, as the overall replication rate of the virus would be determined by the concentration of a particular fatty acid in the different cell strains tested.

In an attempt to validate the potential protease activity and if the bound zinc plays a role in viral replication, we attempted to implement mutations to the zinc coordinating residues in a HEV replicon, based off of the well-studied pSK-HEV2 construct (GenBank: AF444002.1) (16, 35). Wild type and mutant replicons, containing the previously described replication-abolishing GDD to GAD mutation in the conserved motif of the viral RNA-dependent-RNA polymerase (35), were synthesized with a NanoLuc-PEST reporter positioned in frame with the ORF2, and sequence verified. Studies with the pSK-HEV2 replicon typically require the use of the Huh7-S10-3 liver cell line (36), which we were unable to secure access to, and unfortunately all other cell lines tested in-house were unable to support HEV replication. Although we could not execute studies with the HEV replicon, these experiments are vital to gain a more detailed understanding of the HEV life cycle and we are more than willing to provide our replicons to any research group who would be able to follow up with these studies.

Region 510 – 696 was identified to sit between two predicted regions of disorder, so it is plausible that the crystal structure constitutes the complete domain. To confirm this, additional constructs were produced that extended the HEV^510-691^ at both the N- and C- termini and analyzed. NMR studies determined that additional residues added at the C-terminus were disordered, which was consistent with both the secondary structure prediction and the assessment that these residues were part of the disordered ‘hinge’ region. Incremental addition of residues at the N-terminus caused a significant reduction in protein expression, with no protein expression observed with HEV^403-691^ or other constructs extended at the N-terminus. Addition of 70 amino acids to the N-terminus (HEV^440-691^) of the HEV^510-691^ did not seem to have an adverse effect on protein expression, but resulted in the production of a protein oligomer, which was brown in color due to the presence of bound iron. The 70 additional N-terminal residues added contain six cysteine residues, some of which had previously been implicated as potential catalytic residues for the HEV protease (25). Contrary to this report, we would like to propose that these cysteine residues are involved in the formation of an iron-sulfur cluster. This is consistent with observations of iron-sulfur clusters in other hepatitis viruses (38).

As HEV continues to infect a significant number of individuals each year, a greater understanding of the viral biology is required so dedicated antiviral therapeutics can be developed. Until now, the presence of a protease or other non-structural protein between the predicted Y-Domain and hyper-variable region in ORF1 was based upon comparison to other positive-sense RNA viruses and mutagenesis data of residues predicted to be involved with a proteolytic mechanism. Although published data is ambiguous about the presence of a protease in ORF1, the results published in this manuscript confirm the presence of a non-structural protein in the HEV ORF1 between residues 510 and 691. Having characterized the domain, both structurally and biophysically, it is clear the protein has structural homology to a fatty acid binding domain, contains a non-characteristic catalytic zinc-binding motif and could potentially act as a zinc metalloprotease. While we have shown *in vitro* that the identified zinc coordinating residues are essential for metal coordination, we have been unable to confirm the importance of the zinc-binding motif in viral replication. Although we were unable to perform these studies, it is our hope that another research group will follow-up on characterizing the identified domain *in vivo*. It is our belief that the data presented here has provided a significant advancement in the understanding and characterization of the HEV non-structural proteins and we hope that it will form the building blocks for a more detailed understanding of HEV biology.

## Materials and Methods

### Cloning of HEV Constructs

The complete HEV ORF1 (Uniprot ID: P33424) was synthesized as an E. coli codon optimized gene from Synthetic Genomics Inc. Constructs were amplified from the complete optimized ORF1 sequence with primers containing 20 bp of homology (Table 3) to the pET24a plasmid that had been modified to contain a His8-MBP-TEV sequence between the Nde1 and BamH1 restriction sites. The PCR product was analyzed by gel electrophoresis using a precast 1 % agarose gel (Invitrogen), and the PCR products of the desired length were purified using a QIA quick Gel Extraction kit (Qiagen). The purified PCR product was assembled into the vector using Gibson Assembly (NEB Gibson Assembly 2x Master Mix), and transformed into E. cloni 10G chemically competent cells (Lucigen, Middleton, WI). Colonies resistant to kanamycin (Kan) were grown in LB medium, and the plasmid was isolated via Miniprep (Qiagen) prior to being sequence verified using forward and reverse primers. Site directed mutagenesis of the HEV^510-696^ was performed using primers detailed in Table 1 and the Q5 Site-directed Mutagenesis Kit (NEB).

**Table 3:**
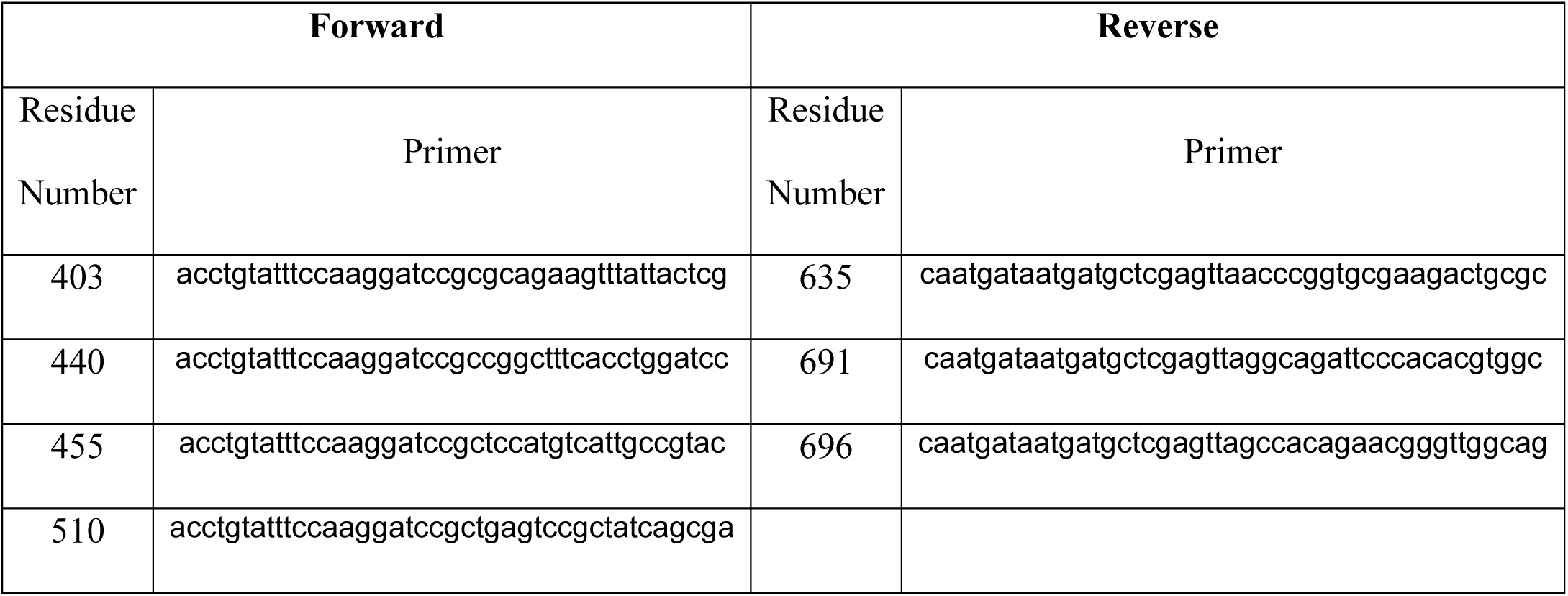
Primers used to clone all HEV constructs.

### Growth and Expression of HEV MBP-fusion Proteins

*E. coli* BL21(DE3) chemically competent cells (NEB product # C2530H) were transformed with 100 ng of pET24a His8–MBP plasmid (Thermo Fisher Scientific) encoding the desired HEV constructs and incubated overnight at 37 °C on LB Agar plates containing kanamycin (50 μg/mL). 50 mL of nutrient rich LB medium containing kanamycin (50 μg/mL) was inoculated with a single colony and grown overnight at 37 °C with agitation at 250 rpm. Using the overnight starter culture, 1 L of Terrific Broth (TB) supplemented with 50 mM 3-(N-morpholino) propanesulfonic acid (MOPS) pH 7.5, 1 x Trace Metals, and kanamycin (50 μg/mL) was inoculated to a starting optical density at 600 nm (OD_600_) of 0.1. If labeled protein was required for NMR experiments, M9 minimal medium where 1 g/L of ^15^N-amonium chloride and 3 g/L of protonated ^13^C-glucose were used as the principal nitrogen and carbon sources, respectively was used instead of TB medium. Cells were grown at 37 °Cwith agitation at 250 rpm until an OD_600_ of 0.8 was achieved and at this stage, the temperature of the incubator was reduced to 18 °C and expression of all constructs was induced using 1 mM isopropyl β-D-1-thiogalactopyranoside (IPTG). Cells were left growing at 18 °C for 18 hours and were harvested by centrifugation, washed with phosphate buffered saline (PBS) buffer and stored at −20 °C.

### Purification of HEV Proteins

Purification of all constructs were performed using the same protocol. *E. coli* cells containing the overexpressed HEV proteins were re-suspended in binding buffer (10 mL/g of cell pellet; 50 mM Tris pH 8.0, 300 mM sodium chloride, 25 mM imidazole), containing Roche Protease Inhibitor without ethylenediamine-tetraacetic acid (EDTA) and passed five times through a microfluidics cell homogenizer (Microfluidics modelM-110P cell homogenizer) at 18,000 psi. Cell lysate was centrifuged at 35,000 g for 60 minutes to remove insoluble cell debris and the soluble lysate was loaded onto a 5 mL His Trap HP Ni column (GE Healthcare) at a rate of 1.5 mL/minute. The Ni resin was washed with 100 mL of binding buffer to remove all non-specifically bound protein before the desired MBP-fusion proteins were eluted from the column using 100 mL elution buffer (50 mM Tris pH 8.0, 300 mM sodium chloride, 500 mM imidazole).

All fractions identified to contain the desired protein were combined and placed in a 30 mL dialysis cassette (Thermo Fisher) with 100 units His tagged TEV protease per 1 mg of MBP-fusion protein, and dialyzed against 2 L dialysis buffer (20 mM Tris pH 8.0, 300 mM sodium chloride, 10 mM imidazole) overnight at 4 °C. The protein was further purified using reverse IMAC with the cleaved protein being loaded onto a 5 mL His Trap HP Ni column (GE Healthcare) at a rate of 1.5 mL/minute. The Ni resin was washed with binding buffer until all cleaved protein eluted from the column. All fractions identified to contain the cleaved protein were concentrated to a volume of 5 mL and loaded onto a HiLoad 16/60 Superdex 75 gel filtration column (GE Healthcare) that had been pre-equilibrated with either crystallography buffer (20 mM Bis-Tris pH 7.0, 150 mM NaCl, 1 mM Dithiothreitol (DTT)) or NMR buffer (20 mM Sodium Phosphate pH 7.0, 150 mM NaCl, 1 mM deuterated DTT) at a rate of 1.5 mL/minute. Samples were concentrated to a final concentration of approximately 10 mg/mL for crystallography experiments, or 1 mM for NMR experiments, flash frozen in liquid nitrogen and stored at −80 °C until needed.

### Crystallography

Crystallization screening was performed using sitting drops, where 400 nL of an equal mixture of protein and precipitant (200 nL each) were incubated against 60 μL of precipitant. Screening was run at two temperatures, 4 and 20 °C, and trays were monitored for crystallization events using robotics. Suitable crystals grew at 4 °C using 18 % w/v PEG 3350, 180 mM LiNO_3_ and 10 mM NiCl_2_. Crystals were cryopreserved using precipitant supplemented with 20 % v/v of glycerol for data collection. All X-ray data was collected on a Rigaku FR-E+ SuperBright generator using a Pilatus R 300K CCD.

As no suitable homologous search model existed for molecular replacement and there was only a single methionine in the protein sequence, the crystal structure was solved using single isomorphous replacement with anomalous scattering (SIRAS). A suitable mercury derivative was generated by soaking crystals in precipitant supplemented with 1 mM of thiomersal for 3 hours. The mercury covalently reacts with the protein primarily at the reactive cysteine residue Cys 649. Initial Patterson map interpretation, phasing, and initial structure determination were executed using SOLVE (39). This initial model comprised approximately 2/3^rd^ of the protein sequence and the remainder of the model was manually built into 2Fo-Fc and Fo-Fc maps using COOT (40). All coordinates were refined to convergence using BUSTER and the PHENIX suite of programs (41– 43). The statistics of data used for refinement is reported in Table 1.

### NMR Experiments

All NMR experiments were conducted with 500 μL of sample containing 5% D2O in a 5 mm NMR tube and were recorded at 298 K on a Bruker AVANCE III 600 MHz spectrometer equipped with a 5 mm CP-QCI-F z-gradient probe. 2D [^15^N, ^1^H]-HSQC experiments were acquired with uniform sampling collecting a total of 1024 and 256 points in the direct and indirect dimensions, respectively. 3D HNCA, HNCO, HN(CA)CO, HN(CO)CA and CBCA(CO)NH experiments were acquired for the assignment of backbone atoms using 50 % non-uniform sampling (44), collecting 2048 points in the direct dimension and a total of 3300 points across all indirect dimensions. Non-uniformly sampled datasets were collected using Poisson Gap sampling schemes and processed in Topspin 3.5 using the hmsIST algorithm (45, 46). All data for the assignment of backbone atoms was analyzed in CARA (47) and residues which experienced a change in chemical shift between the different constructs were mapped onto the protein structure in Pymol (48).

### Differential Scanning Fluorimetry

19 μl of Apo HEV construct (5 μM) was incubated with 10X Sypro Orange fluorescent dye in crystallography buffer and 1 μl of additive. The 20 μl reaction was incubated at 25 °C for 5 min before the temperature was gradually increased to 95 °C at a rate of 1.8 °C/min with fluorescence readings being taken every 0.35 °C. Additives were screened using an 8-point 2-fold dilution series of metals with the top concentration analyzed being 1.25 mM. All measurements were performed in replicates of four, with an average standard deviation of 0.27 °C observed between the replicates.

## Acknowledgments

We thank Drs. Andreas Lingel and Donald Ganem for their advice and mentorship during the project. We also thank Barbara Leon and Dr. Tobias Flecken for advice in regards to construct design, protein expression and purification as well as their useful comments and discussion. Finally, we thank Drs. Suzanne Emerson and Dianjun Cao for helpful discussions about the HEV replicon. All authors have no conflicts of interest to declare.

